# XFlow: An algorithm for extracting ion chromatograms

**DOI:** 10.1101/2019.12.27.889253

**Authors:** Mathew Gutierrez, Rob Smith

## Abstract

Mass spectrometry is a fundamental tool for modern proteomics. The increasing availability of mass spectrometry data paired with the increasing sensitivity and fidelity of the instruments necessitates new and more potent analytical methods. To that end, we have created and present XFlow, a feature detection algorithm for extracting ion chromatograms from MS1 LC-MS data. XFlow is a parameter-free procedurally agnostic feature detection algorithm that utilizes the latent properties of ion chromatograms to resolve them from the surrounding noise present in MS1 data. XFlow is designed to function on either profile or centroided data across different resolutions and instruments. This broad applicability lends XFlow strong utility as a one-size-fits-all method for MS1 analysis or target acquisition for MS2. XFlow is written in Java and packaged with JS-MS, an open-source mass spectrometry analysis toolkit.

## Introduction

Mass spectrometry is a popular approach for measuring the sample-bound content and quantity of a variety of classes of molecules across a broad range of applications including pharmaceuticals, forensics, biochemistry, and food science. All applications of mass spectrometry have a common problem: the instrument itself does not provide measurements of molecules nor their identities, but rather produces raw data that must be rendered human-interpretable through the application of data processing algorithms.

According to community perceptions, advancements in software have lagged behind the steady pace of instrumentation advancements.^1^ Unlike other computational science fields (such as genomics) where several foundational computational problems are regarded as solved, most mass spectrometry users feel that significant problems in computational mass spectrometry remain unsolved^1^ despite (in some cases) dozens of published algorithms designed to address them.^2^ Beyond user sentiment, the experimental influence of algorithm selection suggests that the analysis and advancement of computational mass spectrometry algorithms is a valuable pursuit.^3^

Mass spectrometry systems generate datasets that quantify counts of charged particles at specific mass-to-charge (m/z) values. In liquid chromatography-mass spectrometry (LC-MS) systems, these measurements are taken over the time (retention time or RT) required for the molecules to elute from a chromatography column designed to slow or speed the migration of the molecules depending on particular physico-chemical properties such as size, or polarity.

Mapping the raw LC-MS data points to particular classes of molecule (say, a particular peptide at a particular charge state) provides both an accurate count of the relative abundance of that molecule class (through integrating the intensities in those points) and discriminatory information about the identity of the molecule, as the charge state and uncharged mass can be derived through the m/z gap present between isotopic-specific sub-signals (extracted ion chromatograms or XICs) in the molecule’s signal (see Figure 1).

**Fig 1.**
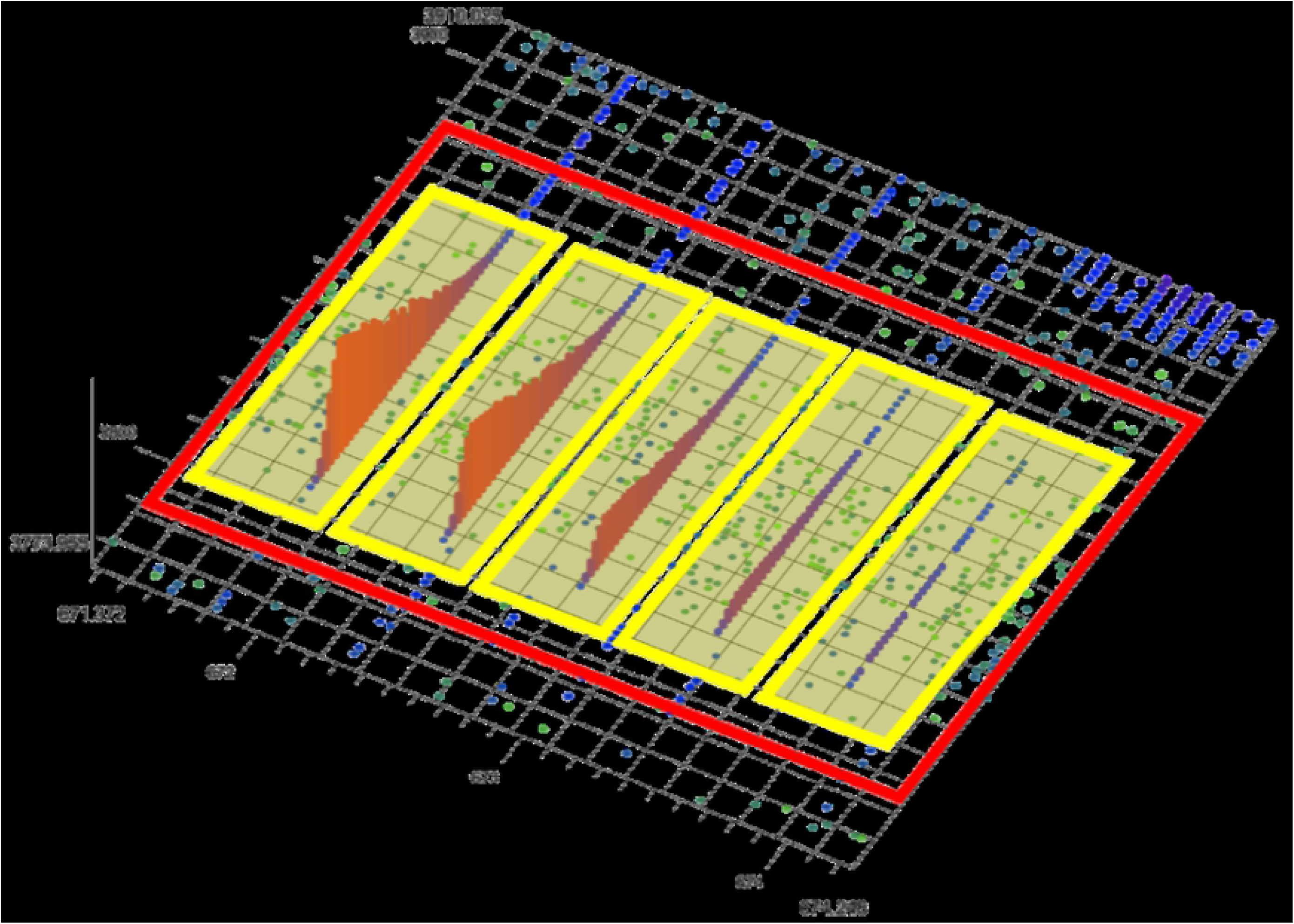
3d Isotopic Envelope. In this figure are five extracted ion chromatograms (XIC) bounded by yellow rectangles. Each XIC is composed of points, each with m/z, RT and intensity (denoted by color and height on z axis). Each XIC is the evidence of an isotope of a specific molecule. The group of five XICs is referred to as an isotopic envelope, or feature, seen bounded by the red rectangle.

Some existing algorithms attempt to resolve the features directly from the point data (e.g. OpenMS FFC^4^). Other algorithms split this process into two steps. First, two step algorithms cluster points into XICs, sometimes called isotopic traces (or features^5^). Second, by clustering XICs into isotopic envelopes (sometimes also called features^5^) (see Figure 1). This two stage approach maximizes the utilization of available information, and serves to reduce the amount of data by allowing a summary of each XIC to be used to find isotopic envelopes instead of cumbersome point data.

This manuscript presents *XFlow*, a novel algorithm for extracting ion chromatograms from LC-MS data. XFlow outperforms existing XIC algorithms evaluated recently on a benchmark human-curated dataset and provides qualitative evidence in support of high-function on alternative datasets. The output of XFlow can be used in conjunction with the XIC clustering algorithm XNet^6^ to map raw data points from an LC-MS run into the signal groups corresponding to particular molecules at particular charge states.

A recent XIC benchmark study^7^ noted that Massifquant^8^, a Kalman filter-based XIC algorithm, performed well against other popular algorithms on a large set of hand-annotated XICs. Massifquant uses a Kalman filter to model XICs as time-series events where the probability of membership of a proximate point in the next scan is a factor of the m/z of previous points in the putative XIC. Massifquant has several drawbacks. It has a very large possible parameter space, takes considerable time to run, and lacks an objective or automatic approach to optimize parameters. Still, it outperformed all other evaluated algorithms on a large human-curated dataset.

Perhaps the principle theoretical advantage of Massifquant is that it attempts to assemble point membership in XICs as a function of the probability of a given point being a member of a given proto-XIC.

XFlow adopts a similar probabilistic approach to constructing XICs. However, it does so with two notable differences. First, the order of the point assembly runs from most intense to least intense instead of from last in retention time to first. Second, the probability function is directly calculable, and does not depend on Kalman filters, which require large matrices that are expensively updated for every point.

XFlow is the first algorithm that leverages intensity order to iteratively construct XICs. Other algorithms use less rich sources of information. Shape filters (for example, matchedFilter^9^ and centWave^10^) tend to degrade at lower intensities and are expensive to optimize for each signal in a run. Massifquant^8^ and MaxQuant^11^ both build XICs scan by scan, though in reverse order. Due to the Gaussian shape of XICs, this guarantees that the least confident information is the most relied upon in both of these algorithms, ensuring suboptimal performance.

The core idea of XFlow is the hypothesis that the most intense points in a mass spectrometry run also have the most accurate m/z measurement. Therefore, intensity of a point can be used as a surrogate for confidence. XFlow leverages this assumption to build XICs starting with the highest intensity points as seeds for putative XICs.

## Methods

XFlow casts ion chromatogram extraction as a clustering problem, where points are clustered into XICs. XFlow creates an initially unlinked graph where each point from an .mzml file (either centroid or profile) becomes a vertex. XFlow groups points together in order of descending intensity by creating links between points nearby in space and intensity. The growing clusters are referred to as “XIC trees”, as each cluster is unique and acyclical in the graph. Due to the relationship between intensity and confidence, each XIC tree is initiated with the highest confidence possible.

Unlike most XIC algorithms, XFlow is designed to be agnostic to instrument and to whether the data is centroided or profile. Unlike any published XIC algorithm, it is also parameter-free. XFlow self-calibrates based on three parameters automatically derived from each run: minimum m/z separation, minimum RT separation, and a two-dimensional grid of the standard deviation of the intensity across the file. The minimum m/z separation between any two points belonging to the same scan is a proxy for the resolution of the machine, and the minimum RT separation between two consecutive scans is a proxy for sampling-rate of the machine. The two-dimensional grid of standard deviation of intensities is used by XFlow to determine a points candidacy by comparison with its neighbors, and not across the file as a whole.

Using the two-dimensional grid of intensity standard deviation, XFlow determines whether each point (p_i_) in the set of all points (**P**) is permitted to enter into consideration and initiate an XIC tree. The justification for this thresholding is two part. The primary consideration of thresholding is to limit the admission of noise into the final output, while the secondary consideration is to reduce the computational burden to only the relevant subset of the data. The study of when and where to apply intensity thresholding is an ongoing and varied topic of research due to the difficulty of avoiding bias, limiting noise inclusion, and maximizing signal inclusion.^12^ For a p_i_ to be considered, its intensity must be at least one standard deviation above the mean for its neighborhood for centroid data, and at least three standard deviations above the mean for profile data. For each p_i_ in consideration, a window of comparison within which to compare nearby points must be constructed. This window of comparison is constructed using the sampling rate and resolution calculated previously to define an arbitrarily large window such that each p_i_ will be compared with all nearby points that could be in the same XIC. The justification for defining such a window is purely in the pursuit of limiting the computation required to just the relevant points. The set of points within this window of comparison will be referred to as **W**. Once the set of points **W** with which to compare to each p_i_ has been obtained, the linking process between p_i_ and all points in **W** begins, and each point (w_j_) in **W** is considered in order of increasing distance from p_i_. For each w_j_ linked to p_i_, the difference between p_i_ and w_j_ scaled by their distance is subtracted from p_i_’s intensity. In this way, the linking process is driven by p_i_’s intensity, with larger intensity values resulting in more links. The formula for the effect on p_i_’s intensity for each link can be seen below in equation 1.

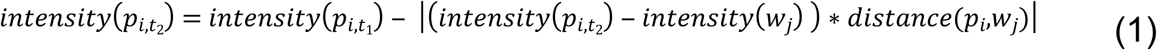

As links are added between p_i_ and points (w_j_) in **W**, XFlow updates the group id of w_j_ to the id of p_i_. This is synonymous with weighted quick union which takes at most M log(N) time for M edges on N objects^13^. If it is the case that p_i_ and a w_j_ belong to the same formative XIC as determined by a shared canonical point, their candidacy is stripped and the next point (w_j+1_) is brought up for consideration. In this way XFlow avoids cycles which can cause problems for resolving subgraphs later. Given that intensity is equivalent to likelihood of participance in an XIC for a point, we can record confidence in any given link as a function of the difference of p_i_‘s intensity before and after linking the point divided by its intensity before linking (see Eq 2). A high confidence point is one that is near in space, and similar in intensity.

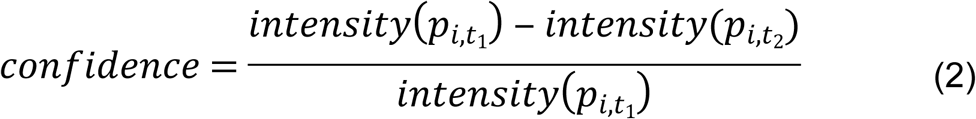

This value is stored such that each link in question has an associated confidence that is a function of the nearness in both intensity and Euclidean distance (given that the difference is scaled by their distance) (see Figure 2). The confidence of each link is used for visualization, but as yet does not affect the composition of the XIC.

**Fig 2.**
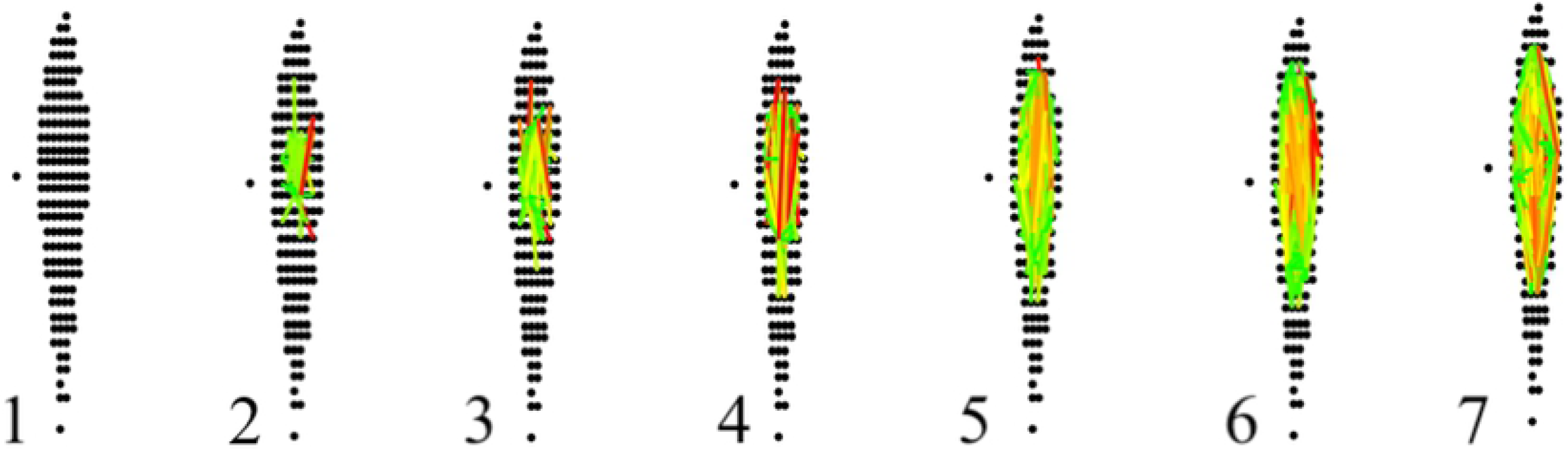
XFlow Link Progression. This figure details the progression of the linking process for a group of points from inception to near completion. In the figure, you can see the completely unlinked group of points that form an XIC (1). Next, the highest intensity point in the group is linked to points with its window of comparison (2). This process continues with the next point, and then the next in descending order of intensity. (3-6). Until all points above the intensity threshold have been recovered (7). Note the correlation between confidence and nearness.

Once XFlow has considered all points (w_j_) in **W** for each p_i_, or exhausted p_i’s_ intensity, it begins the last step, resolving XICs. XFlow resolves XICs by recovering subgraphs created by the points. The subgraphs are elucidated by iterating over all points and adding any XIC with more than five points to the database.

Algorithmic performance is evaluated on a hand-annotated dataset^14^ from a recent study that presented over 57,000 XICs from a public LC-MS dataset^15^ (UPS2). XFlow was compared to the algorithms centWave^10^, matchedFilter^9^, and MZmine2^16^, selected for comparison as equivalent open source algorithms. Accurate evaluation with respect to the hand annotated dataset required point by point comparison. For the chosen algorithms, point data was recovered using the window output that each provided.

For an XIC to be considered appropriately extracted, it must be matched to a corresponding hand annotated XIC. For the purposes of determining accuracy, we will refer to the set of points constituting an XIC produced by the software as **A** while the set of points constituting an XIC produced by hand annotation will be **H**. For an XIC to be considered correctly recovered, the sum of the intensity of the intersection of points between **A** and **H** must constitute greater than fifty percent of the sum of the intensity of the points in the hand annotated XIC (**H**). This fraction of shared intensity will be referred to as *S* (Eq 3).

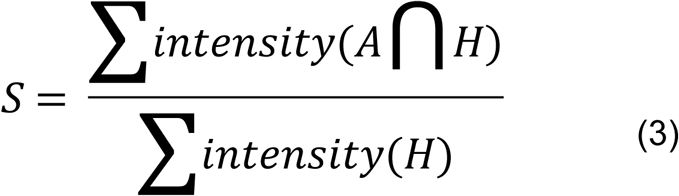

## Results

We compared XFlow to several popular publicly available and functionally equivalent algorithms. XCMS’s centWave^10^ and matchedFilter^9^ algorithms (optimized using Isotopologue Parameter Optimization^17^) and Mzmine2^16^. Due to the difficulty of obtaining verified XIC datasets, quantitative validation of algorithmic results for XFlow, centWave, matchedFilter and MzMine2 are limited to the UPS2 dataset, the only dataset with hand annotated XICs. Five other reference or standard datasets were selected from the PRIDE repository: PXD000790, PXD000792, PXD003236, PXD008952, PXD011194. These additional files were selected in order to provide qualitative information. The RAW files were processed using ProteoWizard’s msConvert^18^ (Version: 3.0.19277-b582d79cd) to create centroided and profile .mzml files using vendor centroiding algorithms. Percent recovery of the hand annotated dataset for each algorithm can be seen in Figure 3.

**Fig 3.**
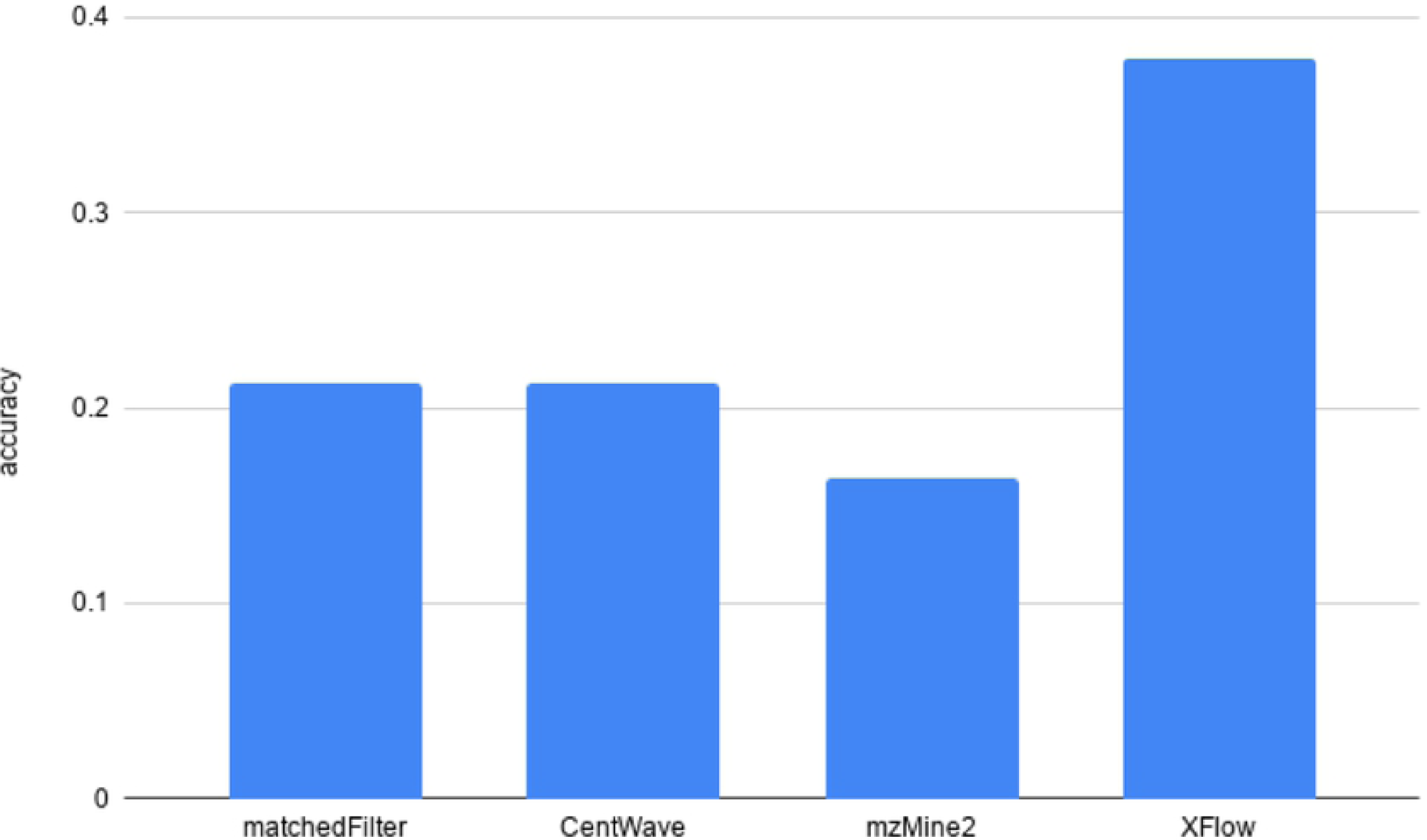
Percent Recovery of Hand Annotated XICs. Percentage of XICs shared between hand annotated dataset, and from the set of XICs output by each algorithm recovered from the UPS2 dataset. Note the relatively low accuracy of XIC recovery across all algorithms, XFlow recovering the most with nearly 40%.

By observing Figure 3, it is apparent that XFlow returns many more XICs from the UPS2 dataset than do the other algorithms chosen. XFlow also manages to recover results closer to hand annotation than the other algorithms. The reason for this is likely the specific method of intensity thresholding XFlow employs, automatically allowing adjustments for each region of the file to be made. Additionally, the difference between signal intensity and noise intensity in the UPS2 dataset is not as great as in other files, and likely the cause of the relatively poor performance across all but XFlow. This relative “flatness” with respect to the other files means that fewer features stand out, providing XNet with its locality depended thresholding an advantage at picking out low intensity signals.

The total number of XICs found in the UPS2 dataset for each algorithm can be seen in Figure 4.

**Fig 4.**
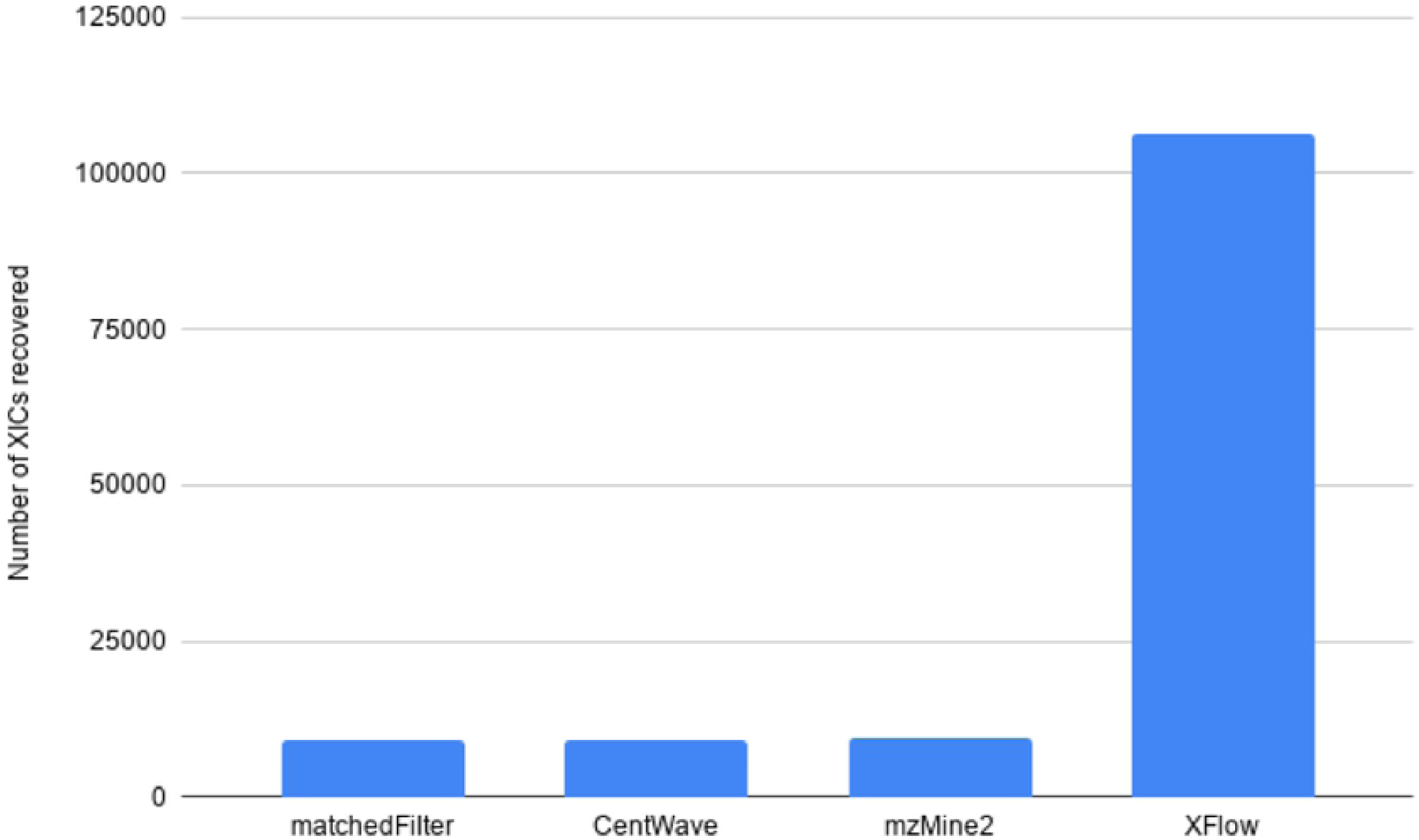
Total Number of XICs Recovered. The total number of XICs in the set of XICs returned from each algorithm from the UPS2 dataset. Note the dramatically increased number of XICs by XFlow compared to existing Algorithms.

The characteristics of a high quality XIC are contiguity along retention time (RT), narrow span along the m/z axis, and a unimodal distribution of intensity along the RT axis. Exemplary XICs from each algorithm on the UPS2 data are shown in Figures 5-8.

**Fig 5.**
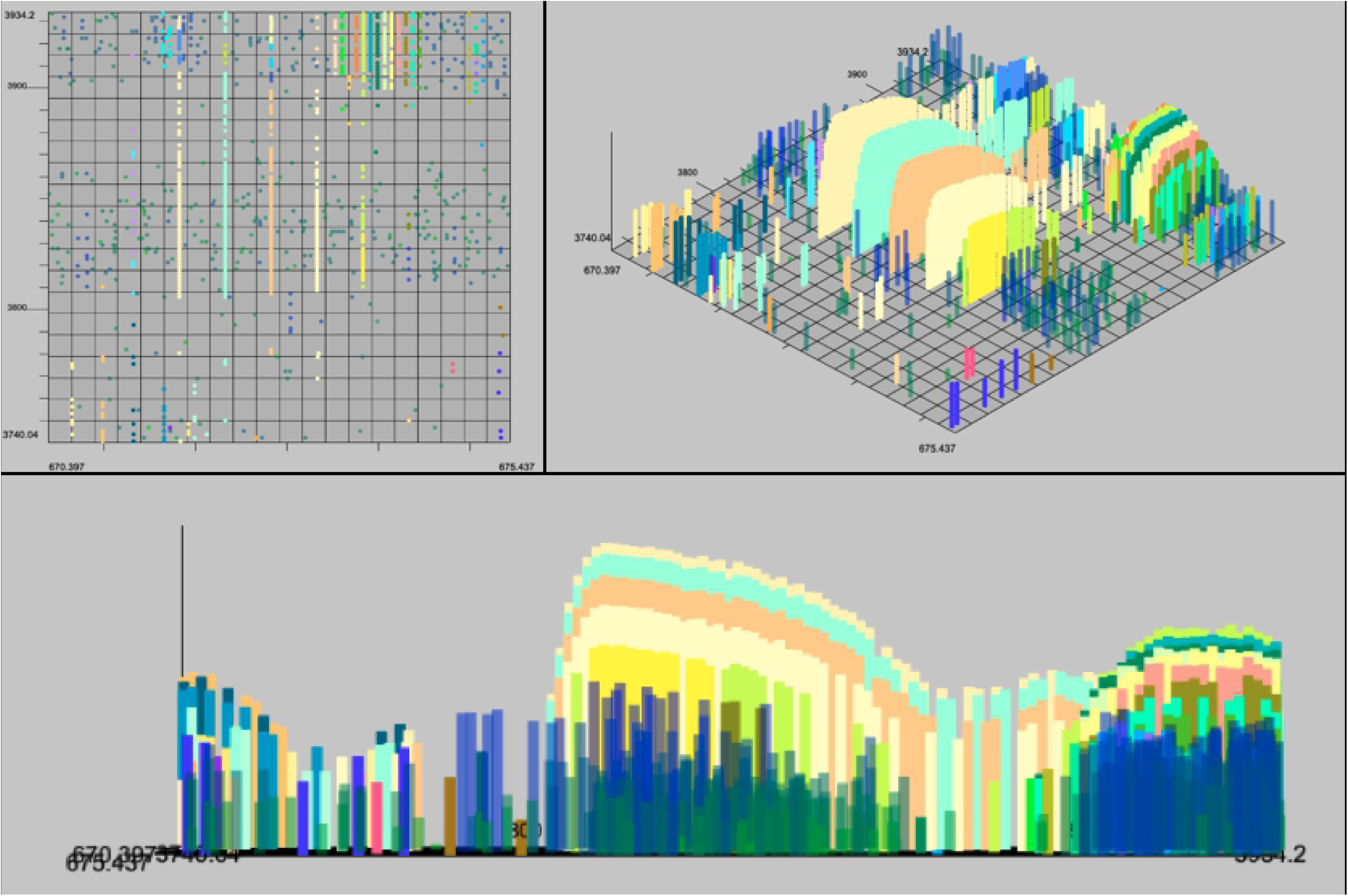
XFlow Results. Selected result of XFlow on the hand annotated data in top-down, 3d and spectral views. All XICs in window are fully formed, and the extent of the XICs in the RT dimension are captured. Note that intensity is not unimodal, and likely a result of another envelope overlapping and adding its intensity to the XICs seen. All algorithms failed to recover these overlapped XICs.

**Fig 6.**
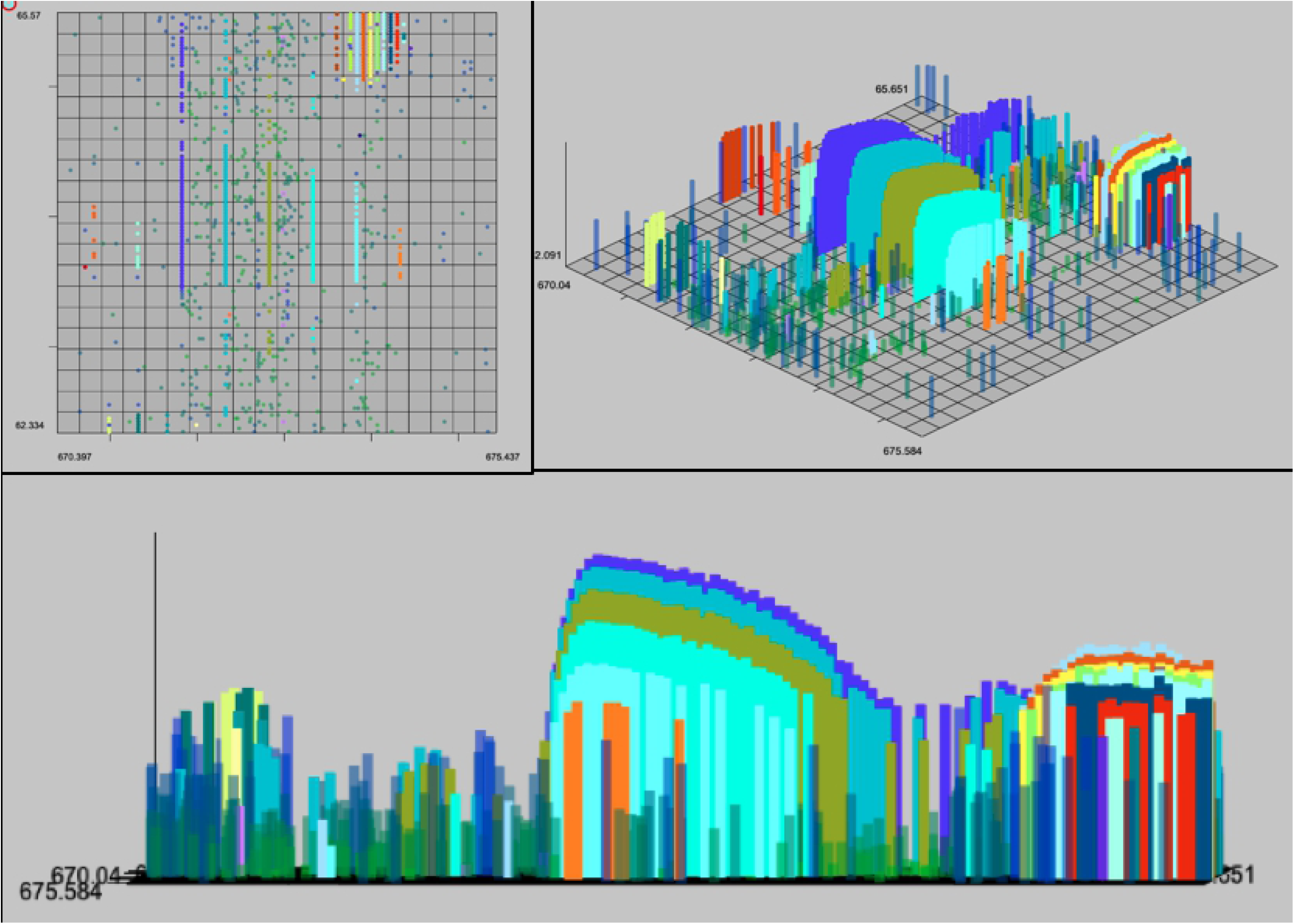
MzMine2 Results. Selected result of MzMine on the hand annotated data in top-down 3d and spectral views. Most XICS in window are fully formed, and XICs extend in the RT dimension to their full extent. Some of the low intensity XICs have been skipped over, as mzMine2 has apparently set the intensity threshold too high to accurately capture all the signals. Like XFLow, MzMine2 has failed to recognize the overlapping signals shown here.

**Fig 7.**
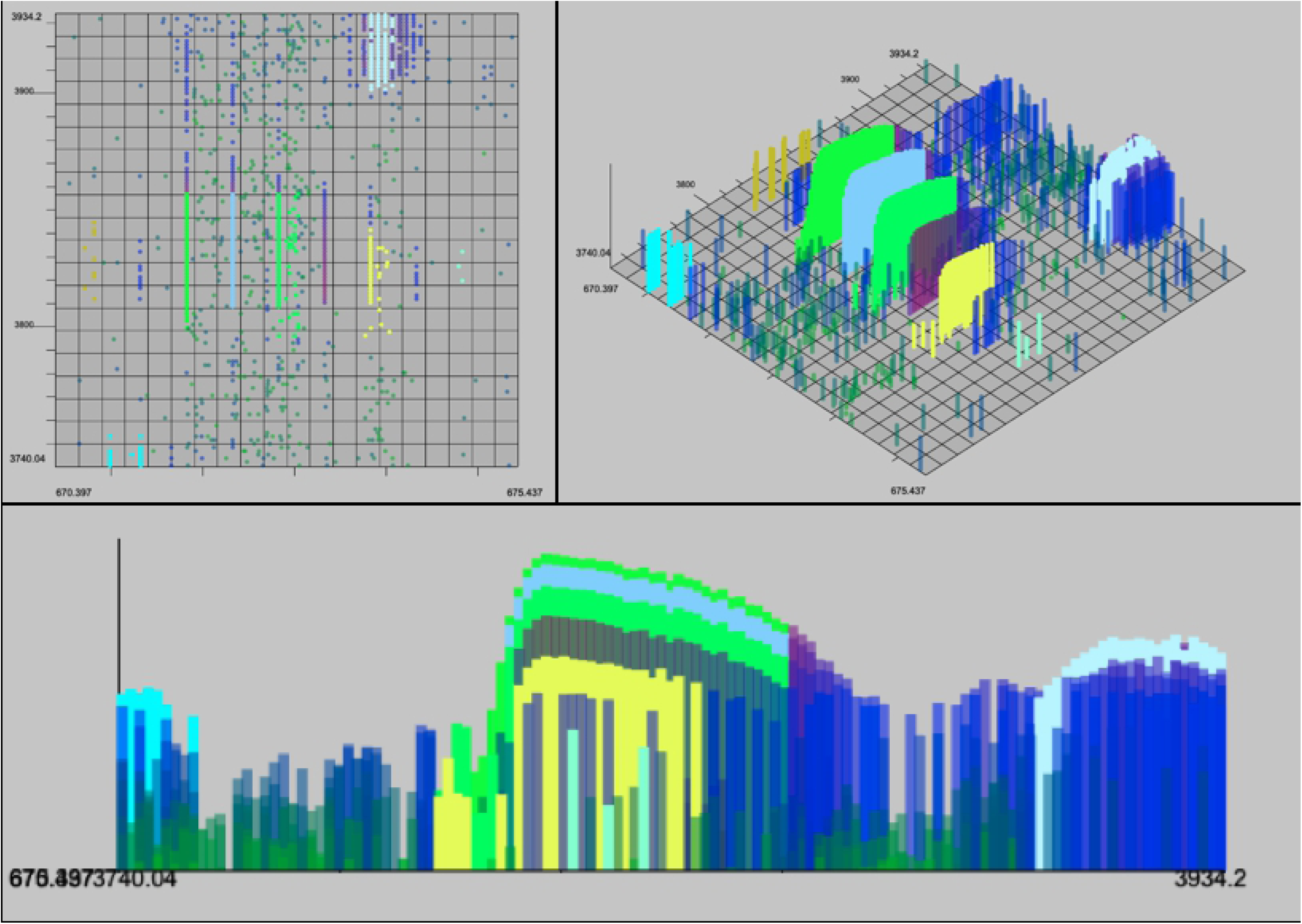
matchedFilter Results. Selected result of matchedFilter on the hand-annotated data in top-down, 3d and spectral views. One XIC in the center has been completely skipped. Several XICs extend too far in the m/z dimension. XICs do not extend fully in the RT dimension.

**Fig 8:**
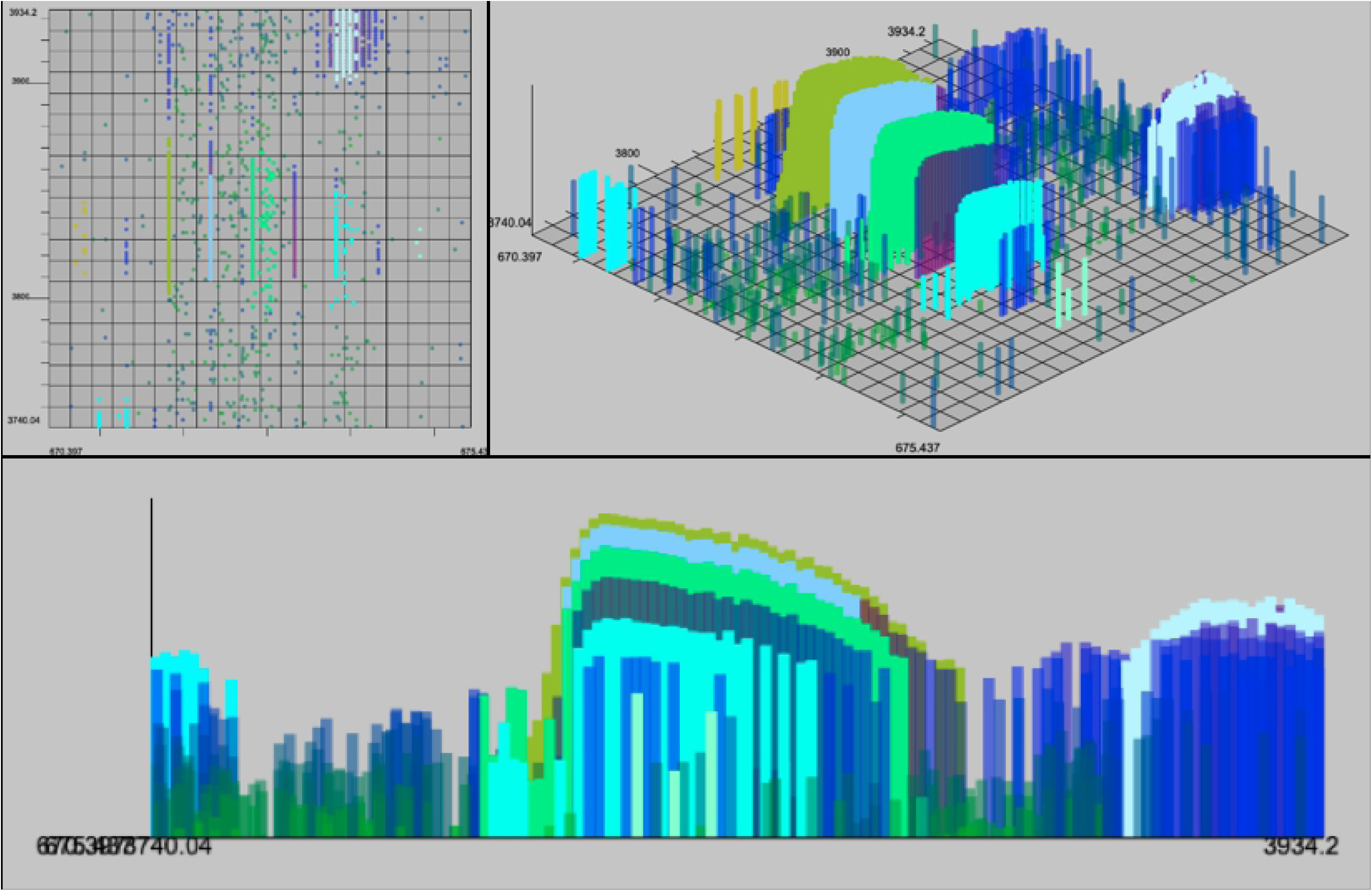
centWave Results. Result example of centWave in top-down, 3d and spectral views. One XIC in the center has been completely skipped. Some XICs extend too far in the m/z dimension as before with matchedFilter. XICs also do not fully extend in RT dimension.

The number of XICs recovered from alternative datasets for both centroid and profile data can be observed in Figures 9 and 10.

**Fig 9.**
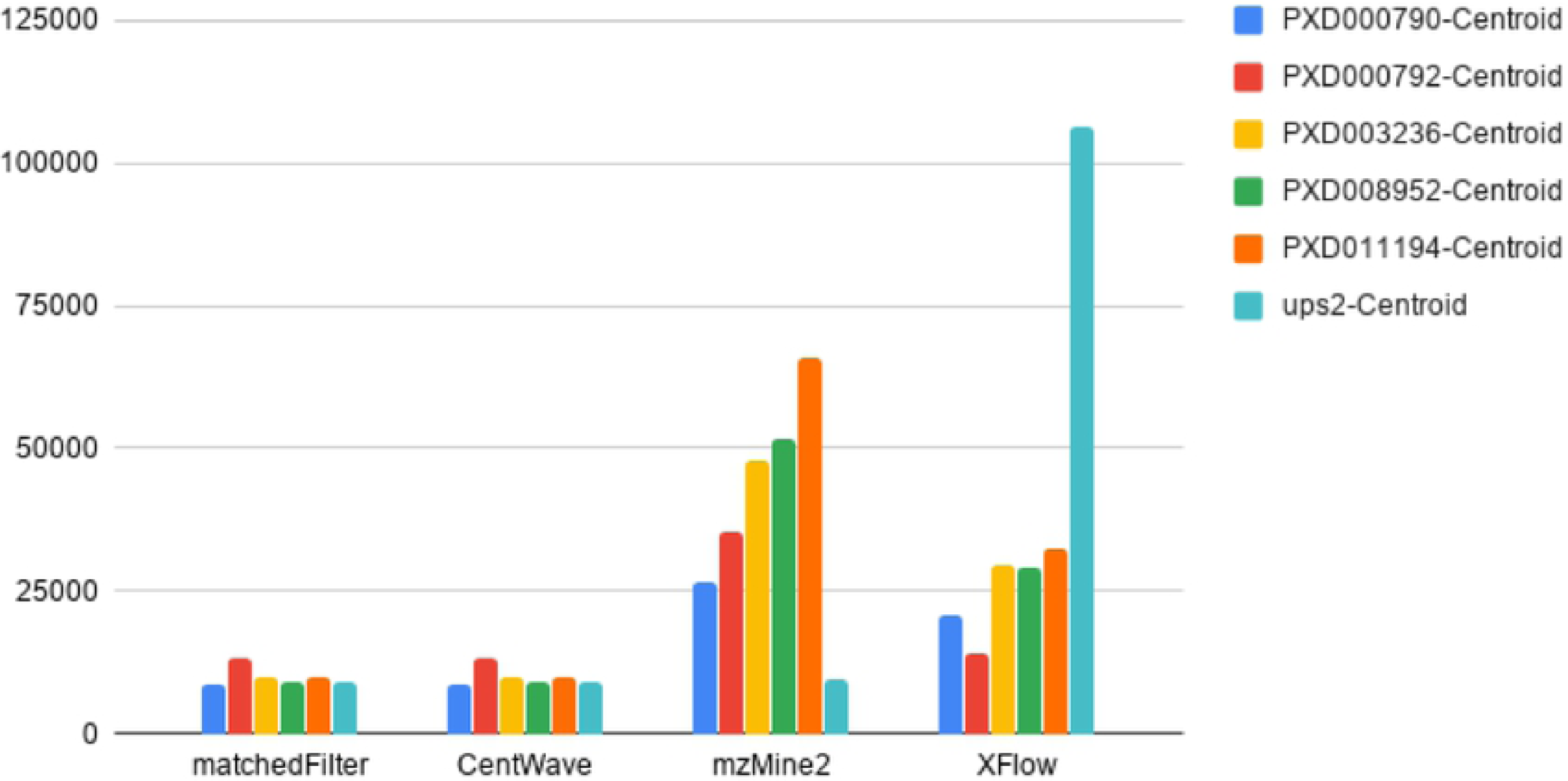
Total Number of XICs for each Centroid File. The number of XICs reported by each algorithm for each centroided dataset. Note the much larger set of XICs returned from XFlow for the UPS2 data set in comparison to other algorithms, and other files.

**Fig 10.**
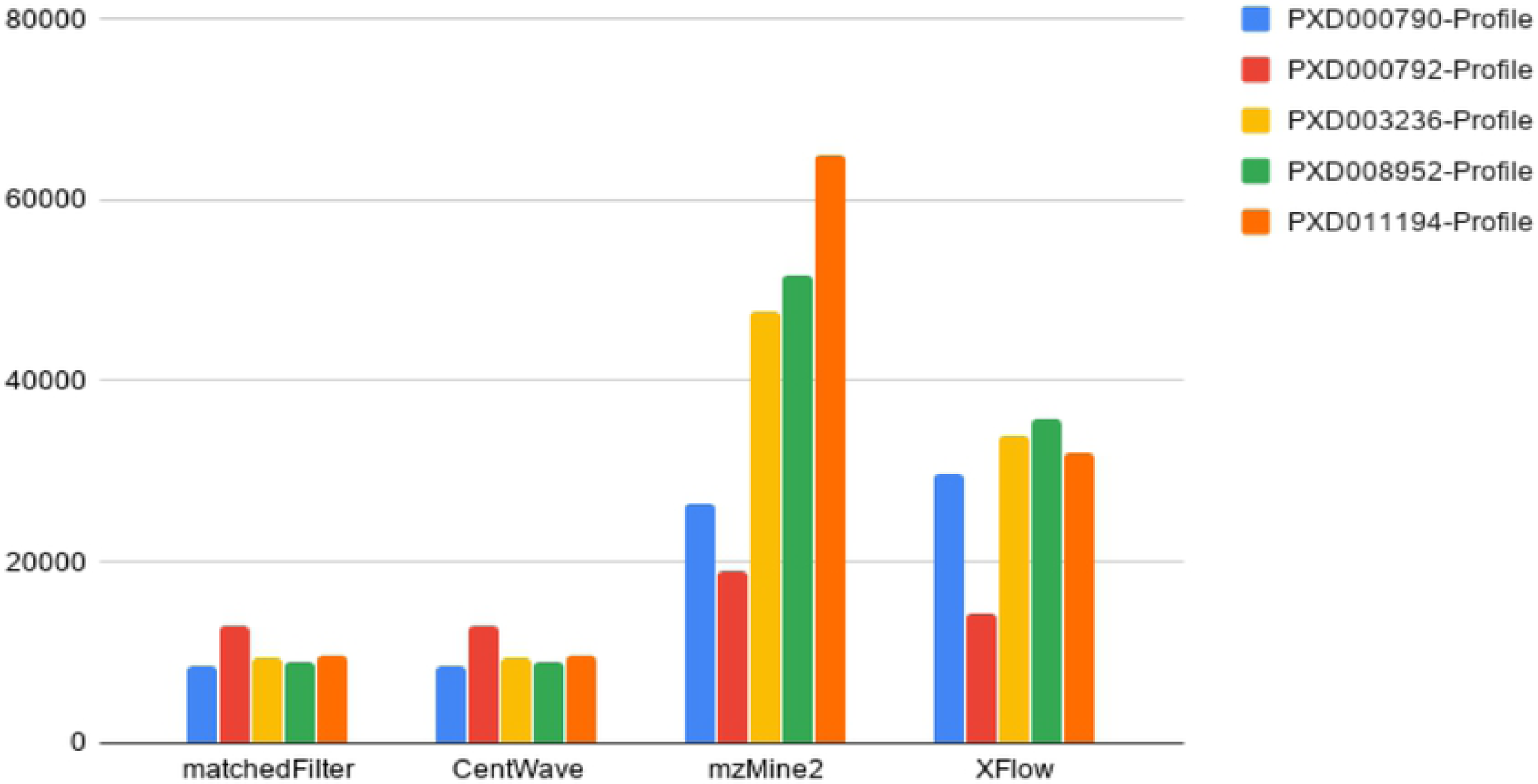
Total Number of XICs for each Profile File. The number of XICs reported by each algorithm for each profile dataset. Note some disparity among mzMine2 and XFlow between centroid (Figure 9) and profile datasets. *Disparity between profile and centroid, while expected, is not desirable. Ideally, the centroid and profile datasets will have the exact same number of signals as they come from the same source. XCMS’s matchedFilter and centWave excelled at recovering the similar numbers of XICs from profile and centroided versions of the datasets*.

XCMS’ matchedFilter and centWave performed similarly in relation to each other. This is likely due to the common origin of the algorithms, and IPO’s optimization strategy. CentWave and matchedFilter also had the most similar results between centroid and profile data (Figures 9-10), also likely attributable to IPOs parameter optimization strategy. The downside of employing IPO is its very lengthy runtime. Further, while centWave and matchedFilter recovered a greater number of the hand annotated XICs, they recovered far fewer XICs from the alternative datasets, qualitatively suggesting that they fail to recover lower intensity signals, an observation that can be verified by analyzing images from PXD011194 dataset to provide qualitative evaluation and comparison between algorithms tested. (See Figures 11-14).

**Fig 11.**
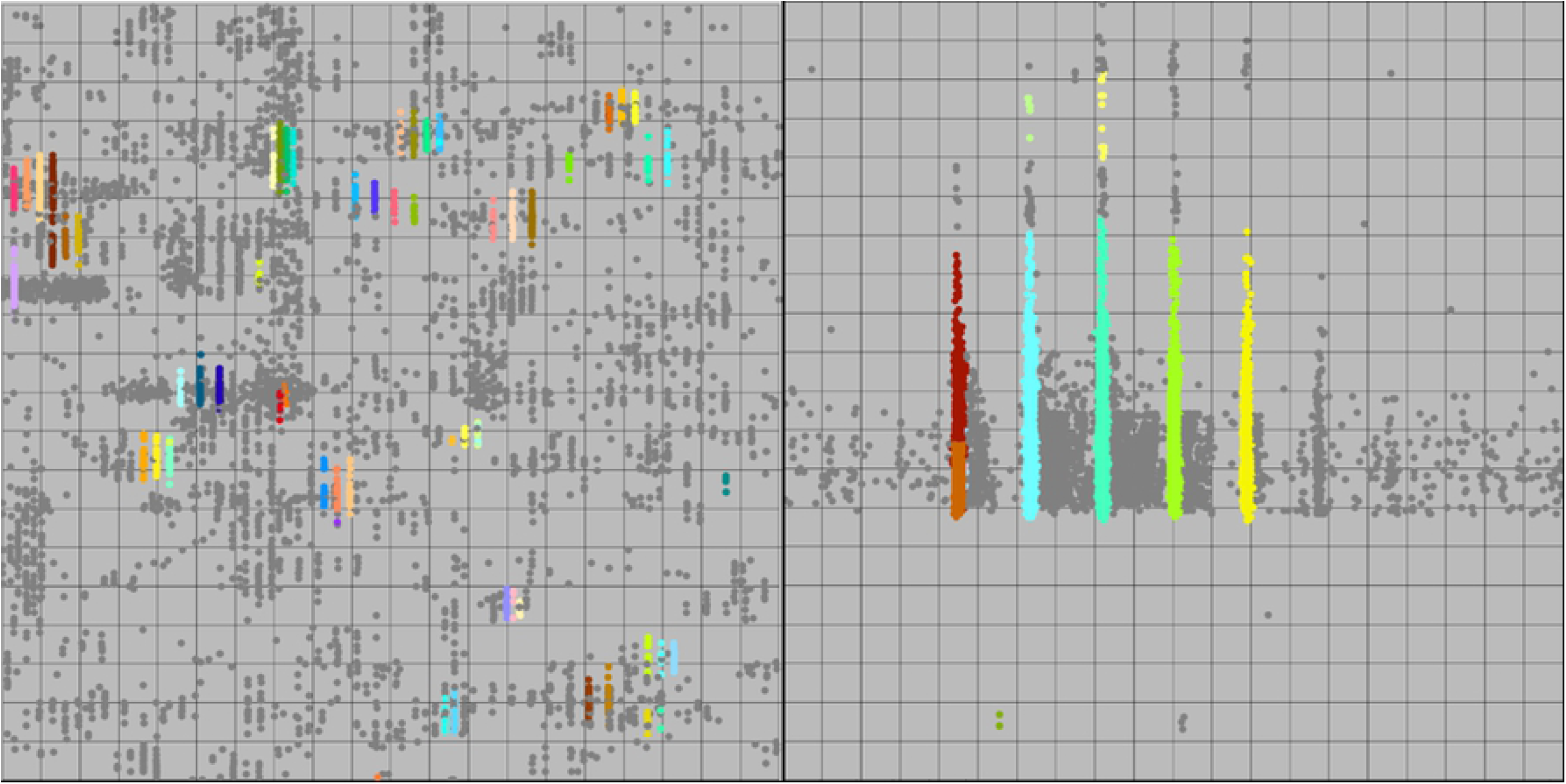
XFlow Qualitative Results. XFlow wide scale (left) and XFlow feature scale (right). It is clear that XFlow has self-adjusted the intensity slightly too high, and as a result is missing clearly present XICS (left). Further, XFlow incorrectly splits the leading XIC in the feature view (right) into two XICs. However, the XICs are well formed, and fully extended, having captured all relevant points.

**Fig 12.**
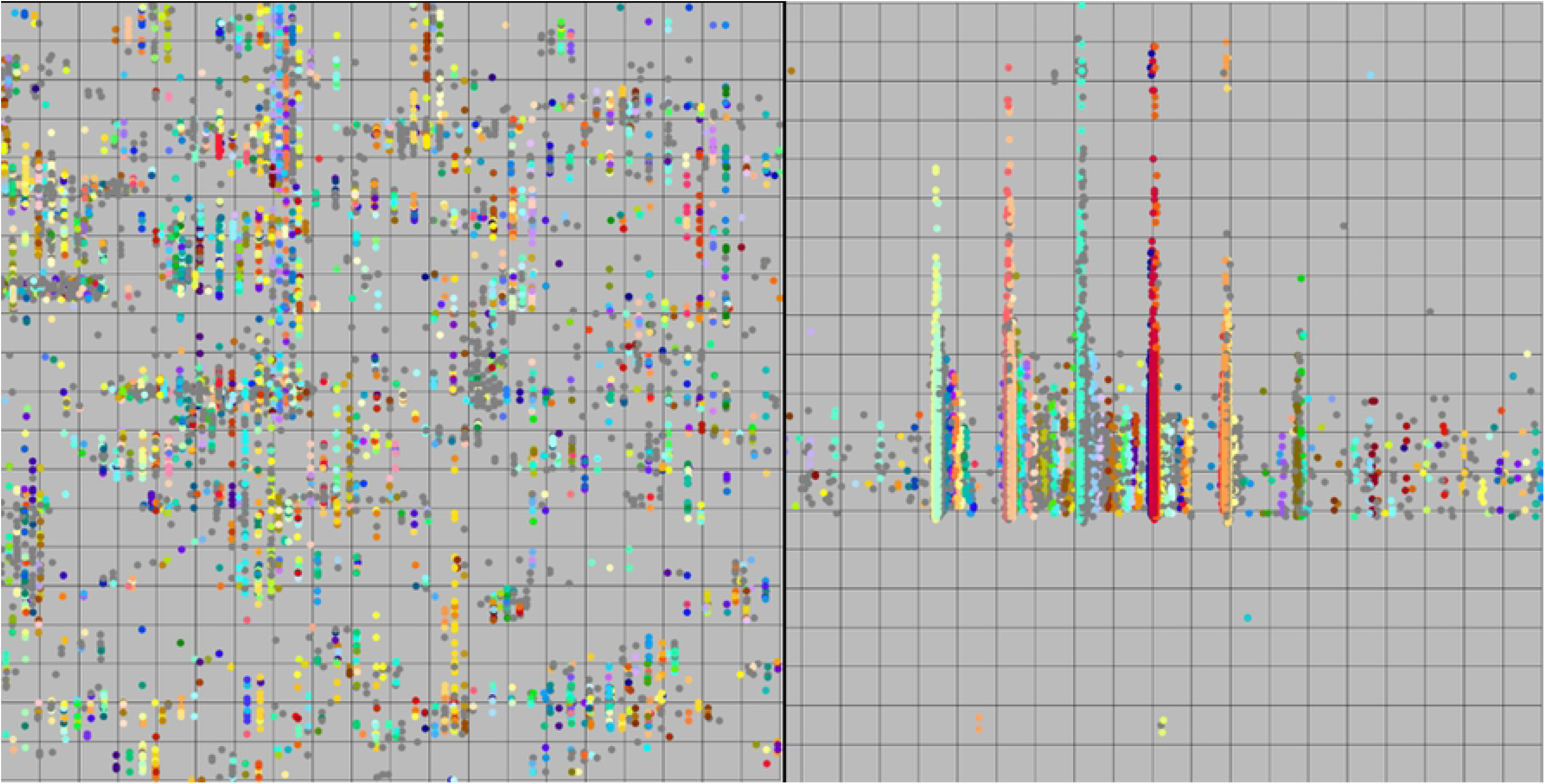
MzMine2 Qualitative Results. MzMine2 wide view (left) and feature view (right). The left view gives the illusion that MzMine2 is selecting for everything not an XIC, however the feature view (right) shows that MzMine2 is finding the XICs, but fails to capture many of the points in those XICs.

**Fig 13.**
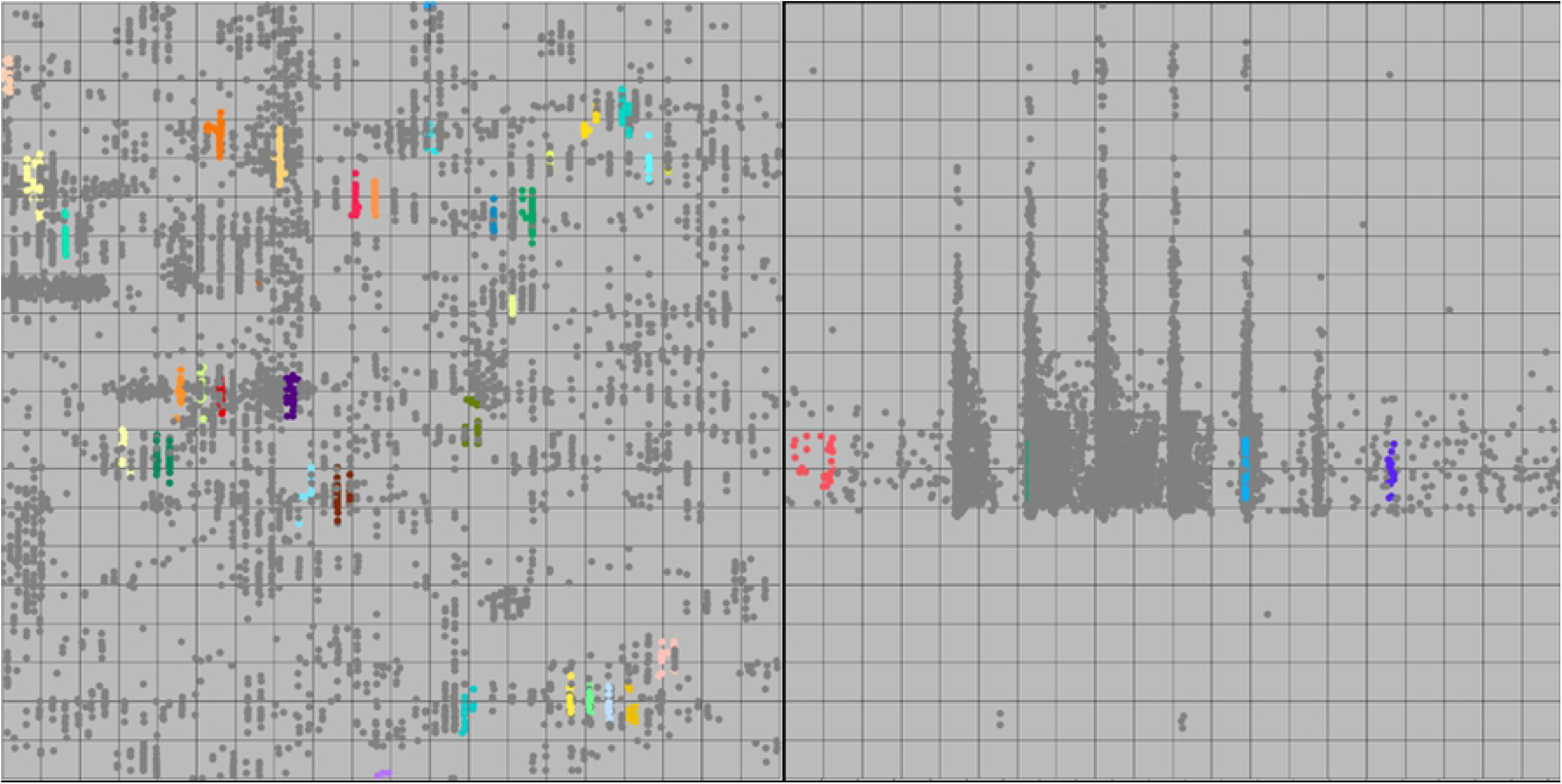
matchedFilter Qualitative Results. matchedFilter wide view (left) and feature view (right). The left view shows that matchedFilter manages to return several of the XICs. MatchedFilter failed to find most of the high intensity points in the feature view (right), but found several of the peaks.

**Fig14.**
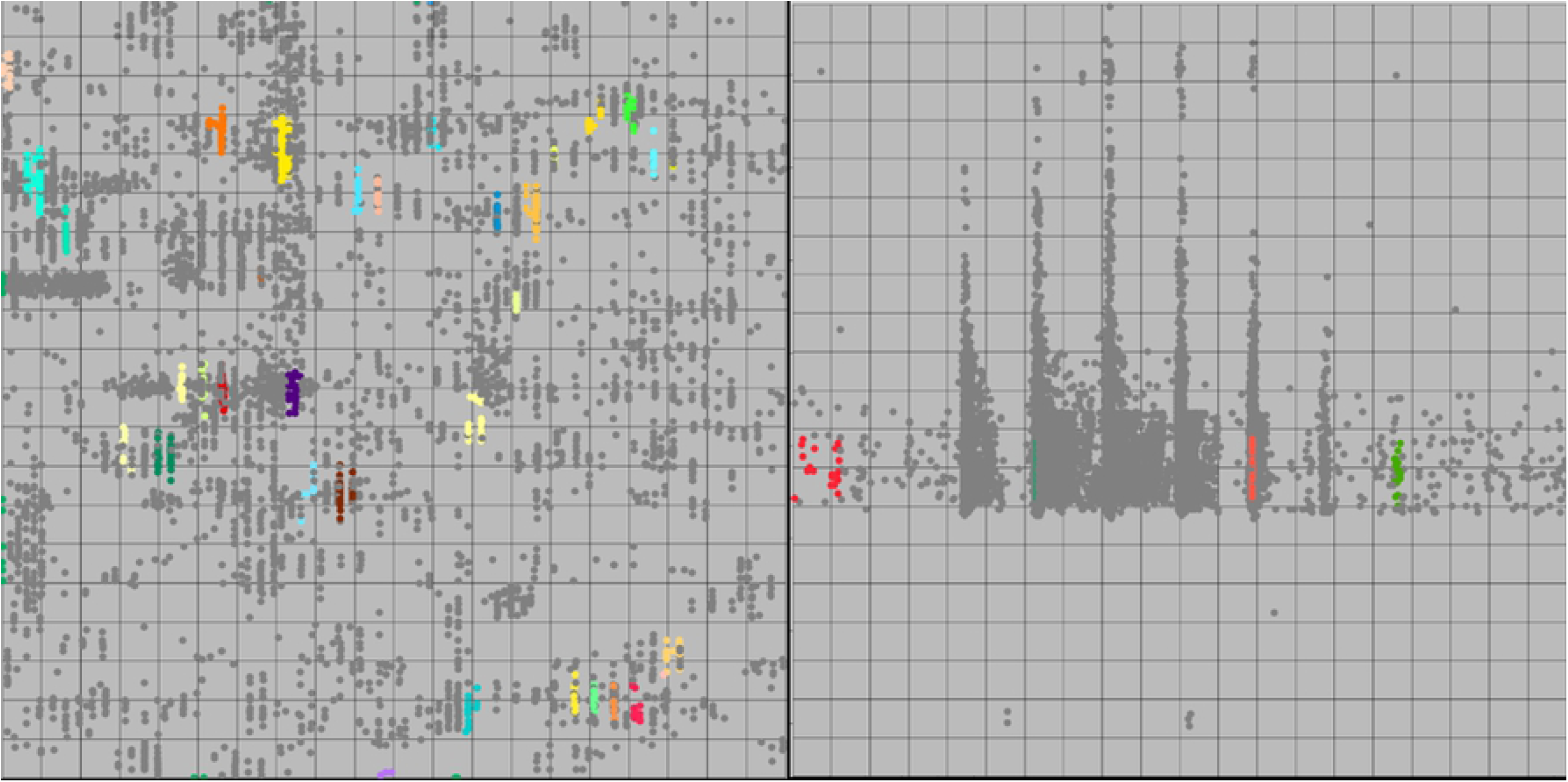
centWave Qualitative Results. centWave wide view (left) and feature view (right). The left view shows that centWave manages to return several of the XICs. Like matchedFilter, centWave failed to find most of the points in the XICs in the feature view (right), but found several peaks.

Additionally, it was observed that centWave and matchedFilter both harbor a tendency to over and under select around regions of interest (Figures 13-14). Additionally, with the prevalence of large datasets, the runtime of these algorithms is vitally important for their continued feasibility in the future. These runtimes can be seen below in Figure 15 (note log scale).

**Fig 15.**
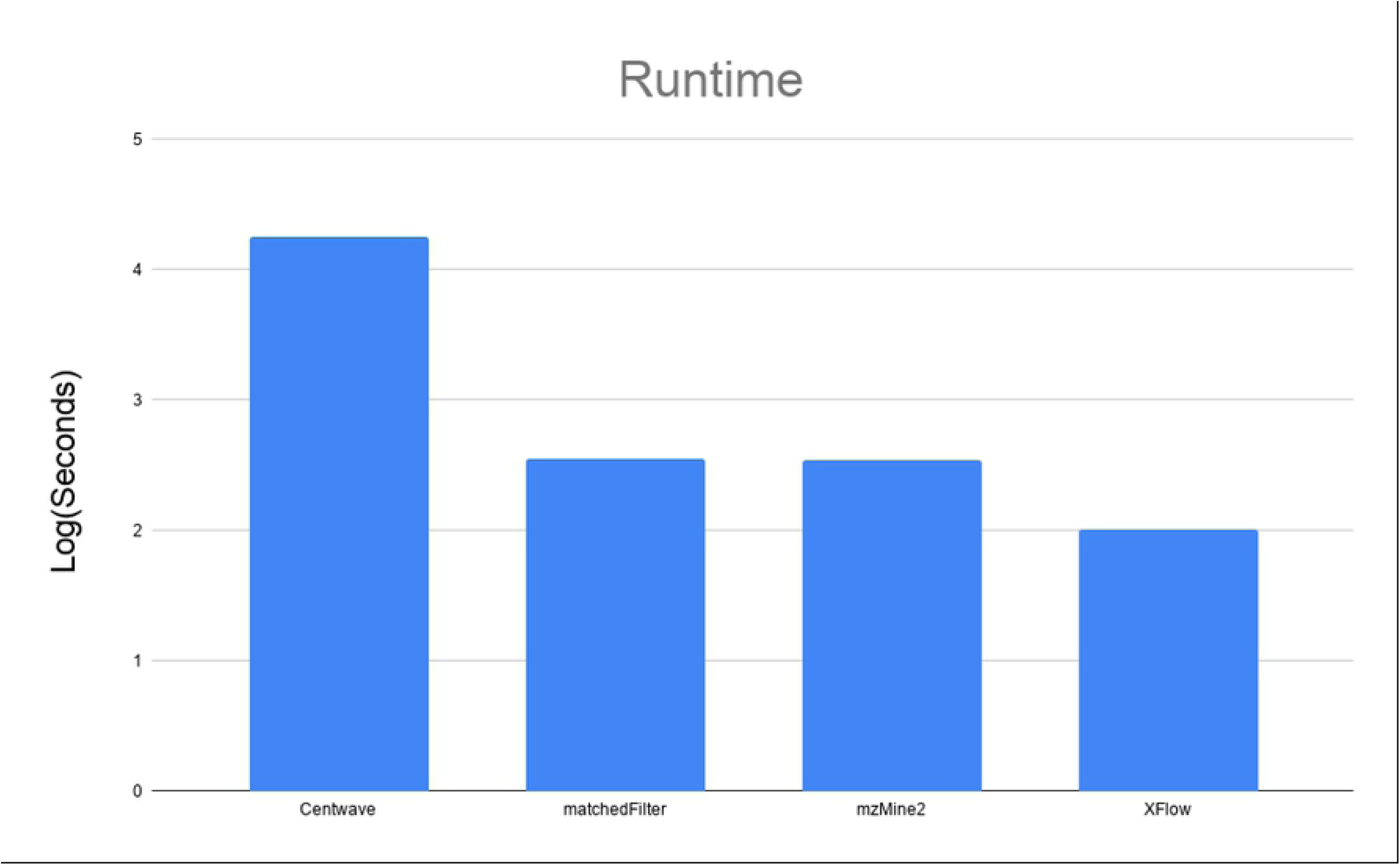
Runtime of Algorithms on PXD000792. This chart shows the log scaled runtimes for each of the algorithms in seconds on the smallest dataset (PXD000792-Centroid). CentWave took the longest at ∼18000 seconds (approximately 5 hours), matchedFilter and MzMine2 took nearly the same at about 5 minutes. XFlow took approximately a minute and a half. Note that the time consuming part of centWave and matchedFilter is the parameter optimization which necessitates repeated trials to obtain optimal results.

## Discussion

The results of this study have brought to light several interesting and key features of the ability of the evaluated algorithms to recover XICs from LC-MS data sets.

MZmine2 was the most permissive of the algorithms tested, resulting in the most XICs recovered compared to the other algorithms (except for XFlow and UPS2) but failed to recover many XICs for the UPS2 dataset, recovering only 18%. Additionally, MZmine2 suffered some disparity between centroided and profile datasets, particularly on PXD000792, a particularly small and low-resolution file. MZmine2 can be qualitatively observed to be too permissive, as many noise points are included as signals (Figure 12), additionally, signal points are often excluded from classification.

Considering the total number of XICs collected for each algorithm with respect to each data file, it is clear here that the UPS2 dataset has interesting qualities in relation to the other datasets (Figure 4). XFlow returned the most hand annotated XICs between any algorithm, and MzMine2 returned far fewer XICs from the UPS2 dataset than any other file, this disparity seems only attributable to something inherent in the dataset itself, likely as mentioned before, the relatively small difference between the signal intensity, and the background noise intensity in the dataset.

The runtime of the algorithms is highly disparate, and the challenges of optimizing a highly parameterized algorithm such as centWave (Even using an automated tool like IPO) is prohibitively time consuming for larger datasets. It is feasible to reuse optimized parameters, but doing so is likely to return suboptimal results. In this way, it is clear that parameterless approaches will excel.

## Conclusion

The size of the datasets, the complexity of the signals, and the noise obfuscation make XIC acquisition from MS1 data extremely challenging. The general method to account for complexity has been to include parameters to increase the scope of an individual algorithm. It was our goal in our lab to reduce complexity, and simplify the experience of conducting MS1 analysis by designing XFlow in a procedurally agnostic way such that it works on a wide variety of MS1 datasets without parameter modification regardless of centroiding or instrument type, a goal that is now accomplished. It’s clear that XFlow excels at signal acquisition for the UPS2 dataset in particular and performs favorably with respect to other algorithms in signal acquisition from alternate datasets (Figures 11-14). Additionally, while qualitative information is gained by comparing results for alternative datasets, it’s impossible to quantitatively evaluate the performance of the algorithms for datasets that do not have a hand annotated version. To this end, developing a database of a variety of hand annotated datasets with which to evaluate algorithms remains a valuable endeavor, in order to provide additional sources of comparison beyond UPS2.

## Author contributions

M.G. wrote XFlow, ran evaluation experiments, curated and interpreted results. R.S. conceived the algorithm and designed and supervised the project. All authors contributed to the manuscript.

## Availability

XFlow is available at github.com/optimusmoose/XFlow with an Apache 2.0 license.Funding

## Materials & correspondence

Should be directed to R.S.

## Supporting Information

XFlow and additional information is available at github.com/optimusmoose/jsms with an MIT license

